# Deepening imaging-based spatial proteomics at high spatial resolution through controlled tissue resizing and in-situ bottom-up mass spectrometry

**DOI:** 10.1101/2025.06.12.659329

**Authors:** Kunyue Deng, Lingpeng Zhan, Li Yi, Jingpeng Zhang, Mingchuan Tang, Tianyi Chang, Gaofeng Ji, Xinyang Shao, Xiang Xu, Xiaoyun Wei, Tianming Zhang, Jiacheng Yao, Jianbin Wang, Guanbo Wang, Yanyi Huang

## Abstract

Understanding the spatial organization of proteins in tissues is essential for elucidating biological function, but current proteomic imaging methods face limitations in resolution, sensitivity, and multiplexing capability. Here, we present IMPACT (Imaging Mass Spectrometry for Proteome Analysis on Compressed Tissue), a hydrogel-based workflow that integrates controlled vertical tissue compression following expansion to enhance protein detection while improving spatial resolution in mass spectrometry imaging (MSI). Through a 6-fold lateral expansion and 100-fold thickness reduction, analytes were concentrated at the surface to improve detection and enable analysis of thick specimen. Coupled with optimized in-gel digestion using hybrid enzymes, IMPACT increased protein identifications 3-fold compared to conventional MSI while maintaining spatial fidelity. We demonstrate IMPACT’s utility in mapping low-abundance proteins and resolving fine anatomical structures. Orthogonal validation with immunofluorescence and integration with spatial transcriptomics further reveal protein-RNA distribution discrepancies, highlighting the need for direct proteomic imaging. IMPACT bridges critical gaps in spatial proteomics by enabling high-resolution, high-sensitivity, and multiplexed untargeted protein imaging, opening new avenues for tissue biology and biomarker discovery.

## INTRODUCTION

Understanding the spatial organization of proteins within biological systems is fundamental to deciphering cellular processes, from signaling pathways to metabolic regulation^1, 2^. While numerous techniques have emerged for spatial proteomics, current methods face critical limitations in specific performances when analyzing intact tissues. Optical imaging approaches, while offering exceptional spatial resolution, are fundamentally constrained by spectral overlap, limiting simultaneous detection to typically <10 proteins^3, 4^. Moreover, antibody availability remains a persistent challenge. Mass spectrometry (MS)-based methods overcome the limitation of multiplexing capability through mass-resolved detection, enabling untargeted analysis of proteome-wide molecules simultaneously^5^. Two primary MS-based strategies have emerged for spatial proteomics: (1) laser microdissection coupled with liquid chromatography (LC)-tandem MS (MS/MS), which achieves deep proteome coverage (up to thousands of proteins) ^6, 7^ but suffers from low throughput and limited spatial resolution (governed by dissection precision), and (2) direct mass spectrometry imaging (MSI) that preserves spatial information through in situ analysis.

Current MSI technologies each present unique advantages and constraints. Secondary ion mass spectrometry (SIMS)^8, 9^ achieves nanometer-scale resolution but is destructive for biomolecule integrity and poorly suited for proteins. Desorption electrospray ionization (DESI) ^10-13^ enables ambient analysis of intact peptides and proteins^14^ but faces sensitivity challenges for large biomolecules^15^. Matrix-assisted laser desorption/ionization (MALDI)-MSI^16, 17^ has emerged as the most versatile imaging platform^18^, combining compatibility with intact proteins^19^ and peptides (natural peptides or proteolysis products)^20, 21^, tunable resolution (5-100 µm), and the ability to generate diagnostic fragments via in-source decay^22, 23^. However, conventional MALDI-MSI struggles with fundamental limitations, including restricted sensitivity due to poor analyte utilization in thick (>30 µm) sections caused by inaccessibility of interior analytes and high electrical resistance^24, 25^, as well as spatial resolution constrained by laser optics and matrix crystallization. While targeted approaches using mass-tags or signal amplifiers^26, 27^ can enhance specific protein detection, they remain fundamentally incompatible with untargeted spatial proteomics due to inherent requirement for prior knowledge of target molecules.

Recent advances in hydrogel-based tissue processing^28, 29^, particularly expansion microscopy (ExM) ^30-32^, have demonstrated that physical sample expansion can overcome instrumental resolution limits by effectively reducing equivalent pixel sizes without modifying instrument parameters. This paradigm has been adapted for mass spectrometry-based spatial proteomics through two strategies: (1) LC-MS/MS analysis of microdissected expanded samples^33, 34^, and (2) direct MSI of small molecules and selected proteins^35-37^. However, current implementations face three fundamental challenges: (i) spatial resolution in microdissection approaches remains constrained by dissection precision, (ii) expanded tissue thickness in MSI methods compromises detection, and (iii) analyte dilution during expansion reduces detection efficiency across all modalities. Addressing these limitations is critical for realizing the full potential of expansion-enhanced spatial proteomics.

Here, we present IMPACT (Imaging MS for Proteome Analysis on Compressed Tissue), an integrated workflow that sequentially combines controlled hydrogel-based 3D expansion, vertical compression of specimen, optimized in situ proteolysis and MALDI-MSI. This method simultaneously addresses the major limitations of conventional MSI through 100-fold thickness reduction that enhances sensitivity by increasing surface analyte density while enabling analysis of thick specimen, following resolution enhancement enabled by tissue expansion. Through systematic evaluation using mouse brain tissues, we demonstrate a 3-fold improvement in protein identifications, including low-abundance functional proteins, and 6-fold enhancement in spatial resolution compared to conventional methods. IMPACT’s capabilities are further validated through orthogonal immunofluorescence comparison and spatial correlation with transcriptomic data, establishing a robust platform for multi-omic tissue analysis.

## RESULTS

### IMPACT workflow enhances MSI sensitivity via 3D expansion and vertical compression of tissue

The IMPACT workflow integrates gel-assisted resizing of the tissue section to be imaged and in-situ proteolysis in gel prior to MALDI-based MSI (Figure 1a). The tissue section is embedded in swellable hydrogel, adopted from expansion microscopy (ExM)^34^ and expansion proteomics (ProteomEx)^33^, where proteins in tissue are covalently anchored to the hydrogel matrix via lysine-acrylyl linkages. Upon hydration treatment, the hydrogel embedding tissue undergoes isotropic 3-D expansion (c.a. 6× linear expansion, c.a. 216× volumetric). To assess spatial fidelity during expansion, we performed 360° radial measurements from the centroid to the tissue border before and after expansion, analyzing magnification within polar coordinates. The expansion factor exhibited high uniformity, with most radial expansions falling within ±5% of the mean (5.93 fold). Following expansion, proteins were denatured with SDS, and non-protein molecules (e.g., lipids, nucleic acids) were removed via washing. Regions of interest (ROIs) were excised from the gel slab under coarse visualization by Coomassie brilliant blue (CBB) and transferred to a cationically modified ITO glass slide, ensuring electrostatic immobilization^38^. Upon dehydration, the gel-embedded ROI underwent vertical compression (from 50 µm to <0.5 µm), while lateral deformation was minimized by electrostatic adherence to the slide. This compression enhances protease/matrix accessibility and reduces electrical resistance, improving ionization efficiency^24, 25^. Contour analysis confirmed minimal distortion (radial deviation: 0.93–1.02), in contrast to unmodified slides. After in-gel proteolysis and matrix application, peptides were analyzed via MALDI MS. Adjacent tissue sections underwent identical sample preparation were analyzed by LC-MS/MS for peptide identification.

**Figure 1.**
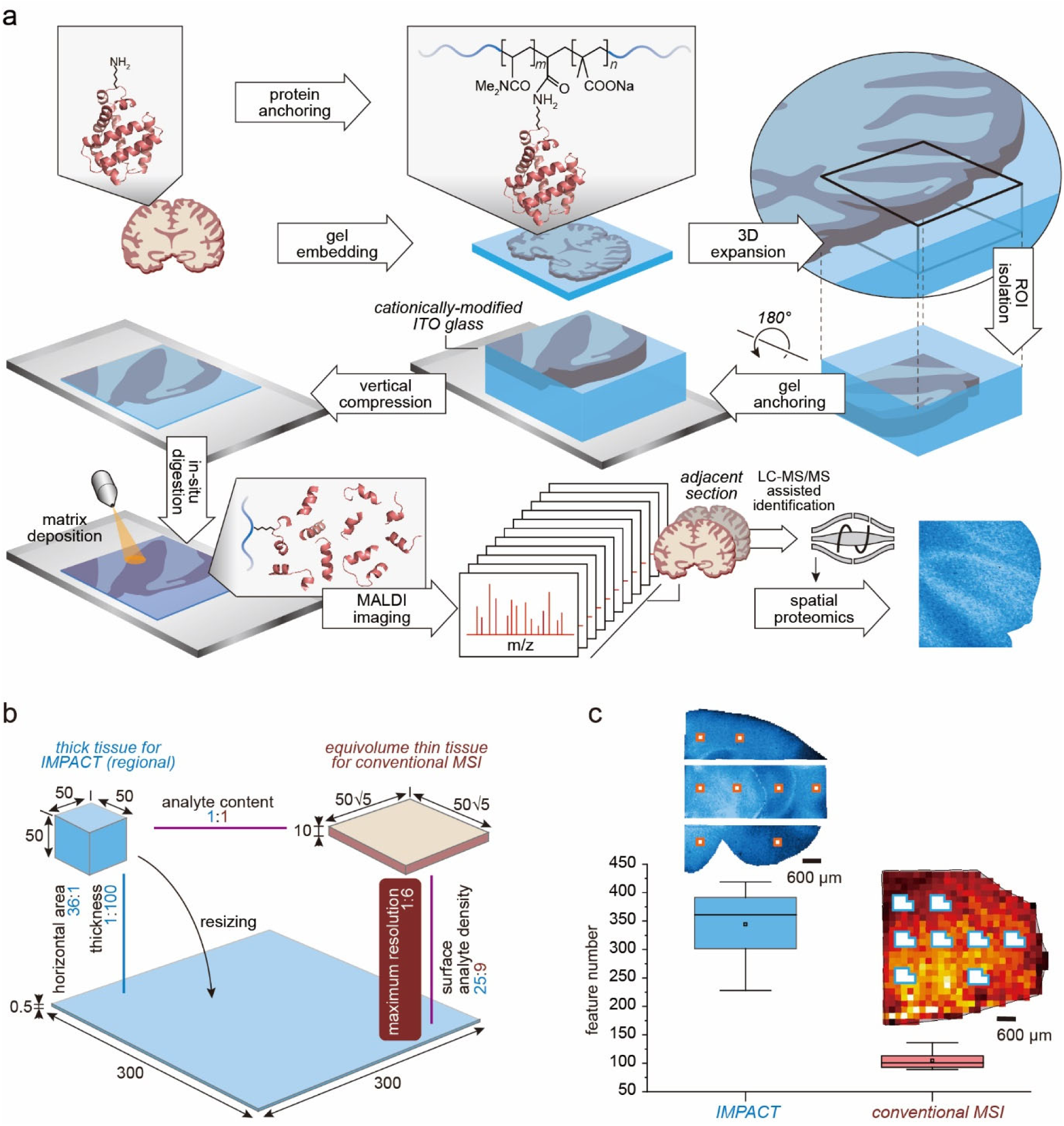
IMPACT workflow and performance advantages. (a) Schematic overview of the IMPACT workflow. (b) Theoretical assessment of IMPACT benefits showing enhanced spatial resolution, tissue thickness reduction, and analyte concentration effects that improve detection sensitivity. (c) Experimental validation of IMPACT’s identification improvement. Box plots display median (central line), mean (central point), and 25th-75th percentiles (box boundaries; whiskers at 1.5 IQR; n=8). Corresponding tissue section images above each plot illustrate the sampling locations and areas analyzed.

MSI spatial resolution depends on both sampling area, which determines the theoretical minimal size of an MSI pixel, and ionizable analyte density, which the practical minimal pixel size that provides adequate analyte ion signal. As illustrated in Figure 1b, IMPACT’s 3D expansion dilutes surface molecule density by 36-fold (square of expansion factor on each linear dimension), but subsequent 100× vertical compression (i.e. concentration) yields a net increase in planar density. Compared to a conventional 10 µm-thick tissue of equivalent volume (containing the same total quantity of analyte molecules), IMPACT offers higher effective resolution (6× improvement for identical laser beam size) and greater surface analyte density (25:9 ratio). In addition, MSI promises improved detection sensitivity due to reduced thickness and enhanced ionization efficiency.

To validate the theoretical advantages of IMPACT, we conducted a systematic comparison of m/z feature detection between IMPACT and conventional MSI. Initial analysis of 50-µm-thick mouse brain tissue sections revealed that IMPACT consistently detected more features than that those reported in previous publication^39^ across varying signal-to-noise (S/N) thresholds. For a more rigorous comparison under controlled conditions, we analyzed adjacent 50-µm (for IMPACT) and 10-µm (for conventional MSI) tissue sections from the same specimen (Figure 1c). To ensure equivalent analyte quantities for direct comparison, we established matched sampling regions with an area ratio of 5:36 (IMPACT vs. conventional). This ratio accounted for the 6-fold linear expansion (36-fold areal expansion) of IMPACT-processed tissue, guaranteeing that each region pair contained comparable initial analyte volumes prior to processing. Across eight such matched region pairs, IMPACT demonstrated a 3.3-fold increase in detected features (344±66 vs 105±16; mean ± SD), validating the sensitivity improvements predicted by theoretical deduction.

### IMPACT enables high-quality MSI of thick tissue sections

Conventional MALDI MSI exhibits significant limitations when analyzing thick tissue sections (>30 µm), primarily due to reduced electrical conductivity that compromises signal acquisition (Figure 2a). The IMPACT workflow overcomes this limitation through vertical tissue compression, enabling robust MSI of thick sections. Figure 2a demonstrates IMPACT’s ability to generate clear spatial patterns for multiple m/z features across the 50-100 µm thickness range, where conventional MSI fails to produce adequate signals for informative imaging. Protein identification via LC-MS/MS analysis of adjacent sections confirmed the biological origin of these features. Furthermore, IMPACT’s enhanced spatial resolution revealed fine anatomical patterns that were undetectable by conventional MSI. Such abilities effectively extend the MSI compatibility from cryosections to vibratome-prepared specimens.

**Figure 2.**
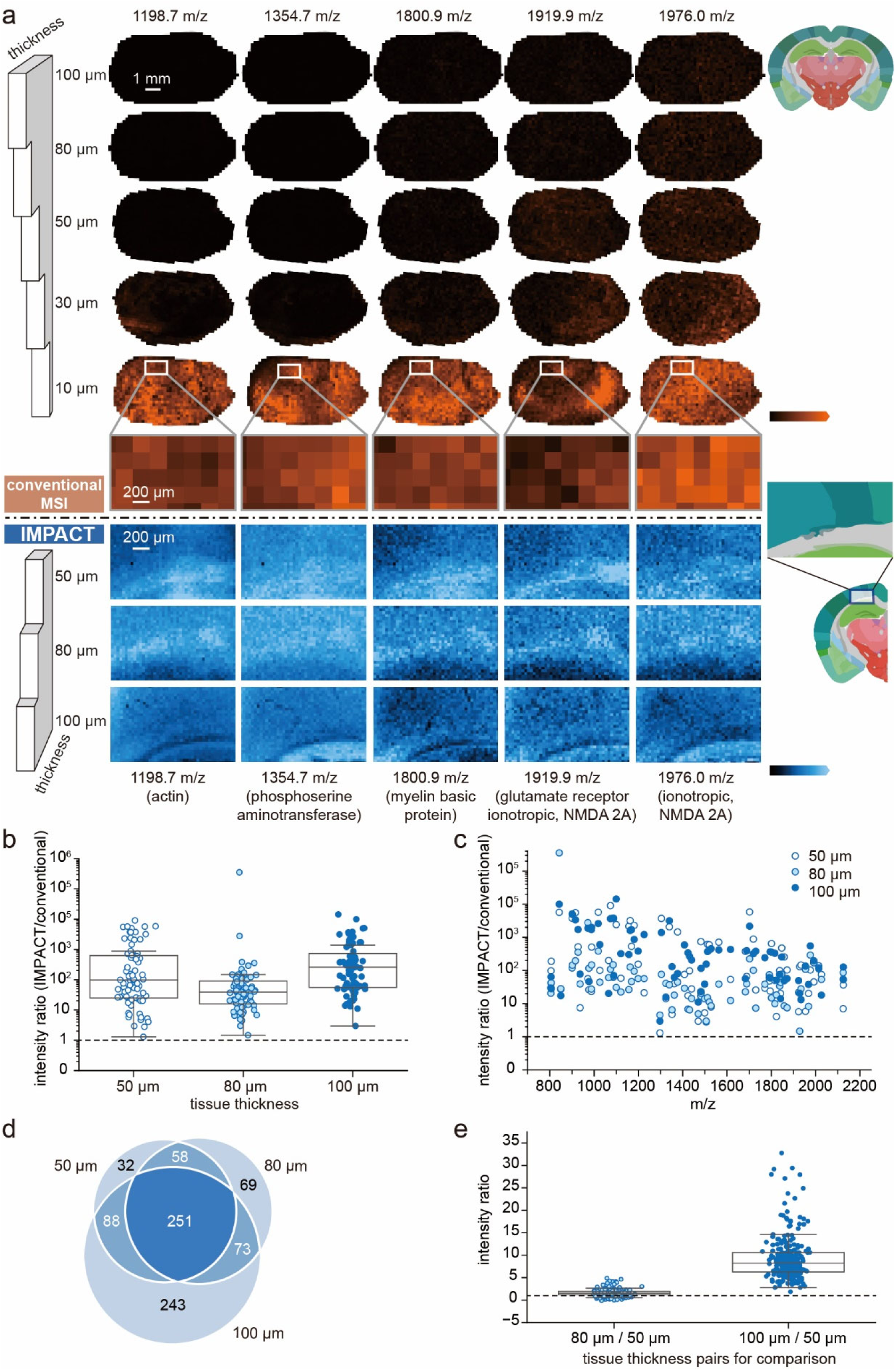
IMPACT enables high-quality imaging of thick tissue sections. (a) Comparative spatial distribution patterns of representative peptides acquired by conventional MSI and IMPACT across tissue sections of varying thicknesses (10-100 µm), with corresponding anatomical references from Allen Brain Atlas (right). (b) Signal intensity ratios of 76 overlapping features detected by both IMPACT and conventional MSI across different section thicknesses (sampling area ≈40 mm²). (c) m/z distribution of features analyzed in panel (b). (d) Feature detection by IMPACT in homogenized mouse brain slices of varying thicknesses (sampling area ≈30 mm²), visualized by Venn diagram. (e) Quantitative comparison of 251 overlapping features in thicker sections (80 and 100 µm) relative to 50 µm reference. Box plot elements: median (line), mean (point), interquartile range (box), and 1.5 IQR whiskers.

Quantitative analysis revealed IMPACT’s substantial sensitivity improvement over conventional MSI (Figure 2b). For the 76 common features detected by both approaches across 50, 80, and 100 µm tissues, IMPACT showed an average signal intensity enhancement of c.a. two orders of magnitude, with consistent performance across all thicknesses. The superior signal quality is visually apparent in the absolute intensity scale spectra. These peptides spanned a mass range of m/z 800-2200, showing a modest detection preference for relatively shorter peptides (Figure 2c). To eliminate potential artifacts from regional protein distribution variations in adjacent sections, we performed parallel experiments using homogenized mouse brain sections. These experiments confirmed that increased thickness in IMPACT-processed specimens led to both higher number of feature detection (Figure 2d) and higher signal intensity (Figure 2e). Method reproducibility was further confirmed through comparative analysis of two separately processed mirror-image ROIs (encompassing cerebrum, brain stem, and fiber tract regions) from bilaterally symmetric positions on the same brain section, which showed highly similar spectral patterns and strong correlation. These findings demonstrate that with IMPACT, greater specimen thickness no longer compromises MSI performance, but instead enhances both detection depth and signal quality.

### Spatial distribution of proteins revealed by IMPACT

The spatial resolution and localization accuracy of imaging markers in MSI can be compromised by analyte diffusion. Since tissue processing prior to MSI involves multiple aqueous treatments, concerns may arise regarding potential diffusion of proteins or their peptide fragments. To quantitatively assess diffusion during the IMPACT workflow, we analyzed the signal intensities of a panel of detected peptides along a horizontal trajectory crossing the tissue border. The overlaid intensity profiles of these peptides revealed a sharp decline at the tissue edge, with less than 5% residual signal in blank regions, indicating minimal diffusion-related interference with the imaging pattern.

To evaluate the imaging performance of IMPACT, we imaged 50 µm-thick mouse brain sections at a spatial resolution of 33 µm (direct sampling pixel size of 200 µm). Due to tissue expansion during gel-assisted resizing and spatial constraints on the glass slide, each section was divided into smaller specimens for high-resolution regional imaging, followed by a reassembly of the regional images (Figure 3a). Peptide identification via LC-MS/MS of adjacent sections, which were subjected to identical gel embedding and sample preparation, enabled attribution of detected masses to their precursor proteins, facilitating protein spatial distribution mapping. The removal of a marginal tissue portion produced a well-defined boundary in the peptide distribution patterns (red arrow, Figure 3a), with no detectable signal beyond this edge, further confirming negligible diffusion within the gel matrix. For proteins such as sodium/potassium-transporting ATPase subunit alpha and myelin basic protein, multiple proteolytic peptides were detected, allowing cross-validation of distribution patterns using distinct imaging markers (Figure 3b). While peptides derived from the same protein exhibited broadly consistent distributions, minor spatial variations in local abundances were observed. These discrepancies may arise from spatially heterogeneous factors, including residual higher-order structures or steric hindrance from background matrix effects that caused site-dependent proteolysis efficiency, and inconsistent signal dilution due to cation adducts (e.g., Na⁺/K⁺) with localized concentration fluctuations. By integrating the distribution patterns of multiple peptides from the same protein, we generated a composite map based on mean abundances, thereby mitigating potential errors introduced by the aforementioned factors (Figure 3b).

**Figure 3.**
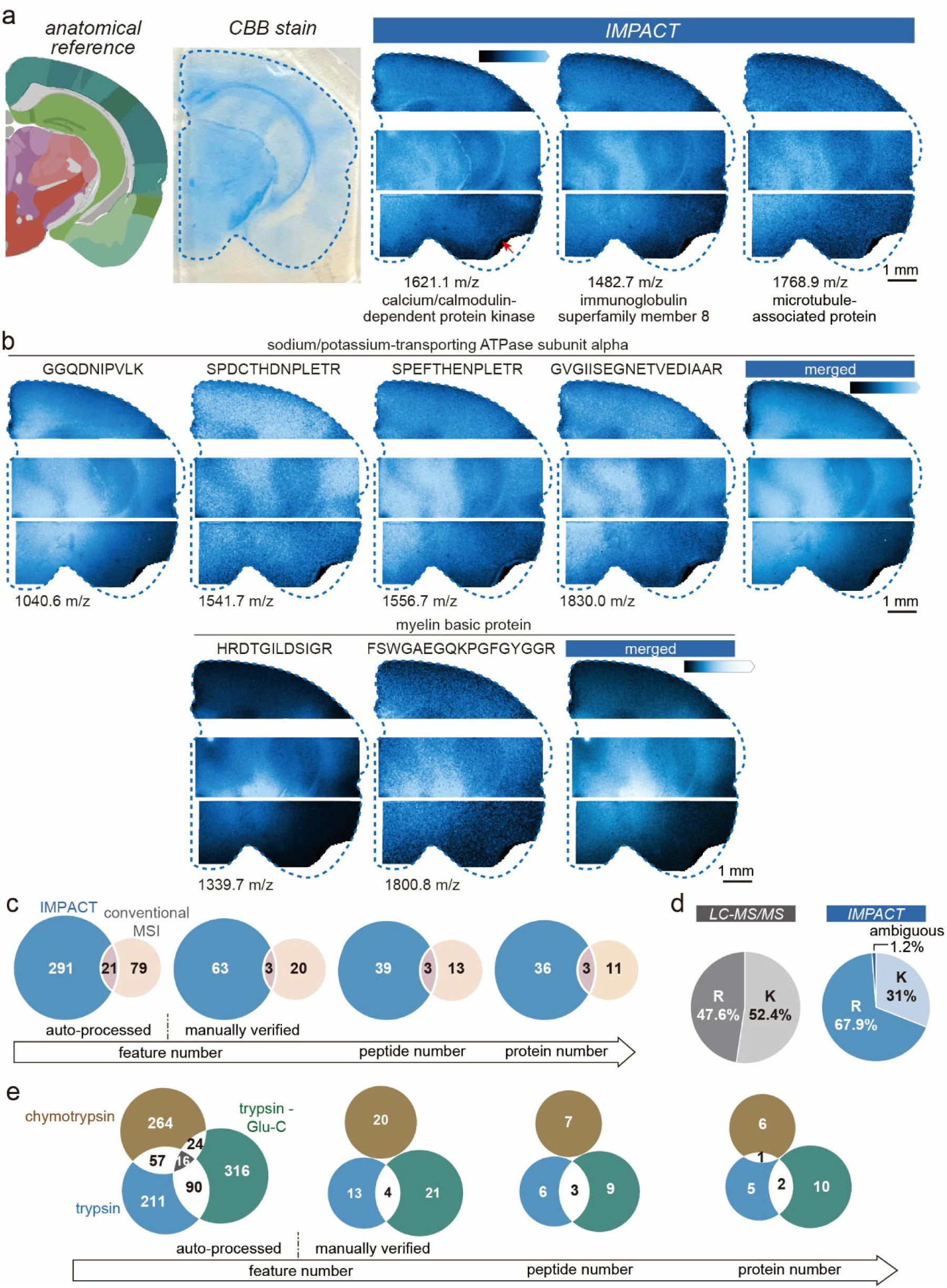
Spatial distribution of proteins in mouse brain sections acquired by IMPACT. (a) Images of representative peptides with protein identifications, shown alongside corresponding anatomical reference (Allen Brain Atlas, left) and CBB stain (middle) of the same section used for imaging. (b) Multi-peptide validation of protein localization: images of individual peptides from sodium/potassium-transporting ATPase subunit alpha and myelin basic protein (left), and merged protein distribution maps generated by signal averaging of corresponding peptides. (c) Number of spectral features, peptides and proteins identified by IMPACT (analyzed section) and conventional MSI (adjacent section) respectively. (d) Proteolytic cleavage site analysis: terminal Arg and Lys distribution of peptides detected by LC-MS/MS (whole mouse brain) and IMPACT (imaged section) respectively. (e) Number of spectral features, peptides and proteins identified for three adjacent tissue sections digested with trypsin, chymotrypsin and trypsin+Glu-C, respectively.

To minimize false discoveries and misidentifications, we performed rigorous manual verification of identification results by correcting deisotoping errors and filtering out misassigned peaks (see Experimental Section for details). Compared to conventional MSI workflows, IMPACT yielded an approximately 3-fold increase in the number of detected features, peptides, and proteins (Figure 3c). Notably, the overlap between identifications from IMPACT and conventional MSI was small, suggesting that each method captures distinct protein populations. This discrepancy likely arises from the gel-anchoring of Lys residues during the IMPACT workflow, which either restricts trypsin cleavage or impedes the release of digestion products for detection. Supporting this hypothesis, we found that while >52% of peptides from whole mouse brain tissue detected using LC-MS/MS terminate with Lys at both termini, only 31% of IMPACT-detected peptides exhibited Lys-terminal ends (Figure 3d). This reduction was further corroborated by analyzing peptides with Lys at both N- and C-termini across tissue sections of varying thicknesses, confirming that gel anchoring blocks a substantial fraction of Lys-containing peptides. To overcome this limitation, we employed complementary proteases with different cleavage specificities, such as chymotrypsin (cleaving after Phe, Tyr, Trp, Leu, and Met) and Glu-C (preferentially cleaving after Glu), either as substitutes for or in combination with trypsin. This strategy increased identification numbers several-fold (Figure 3e), thereby expanding the pool of detectable peptides and enhancing imaging channel diversity.

### IMPACT improves spatial resolution of MSI

To assess the practical advantages of IMPACT in MSI spatial resolution, we performed a systematic comparison with conventional MSI across multiple resolution settings (200 µm, 100 µm, and 50 µm pixel sizes) commonly used for protein imaging (Figure 4). Due to IMPACT’s inherent resolution enhancement, these settings correspond to effective resolutions of 33 µm, 17 µm, and 8 µm, respectively.

**Figure 4.**
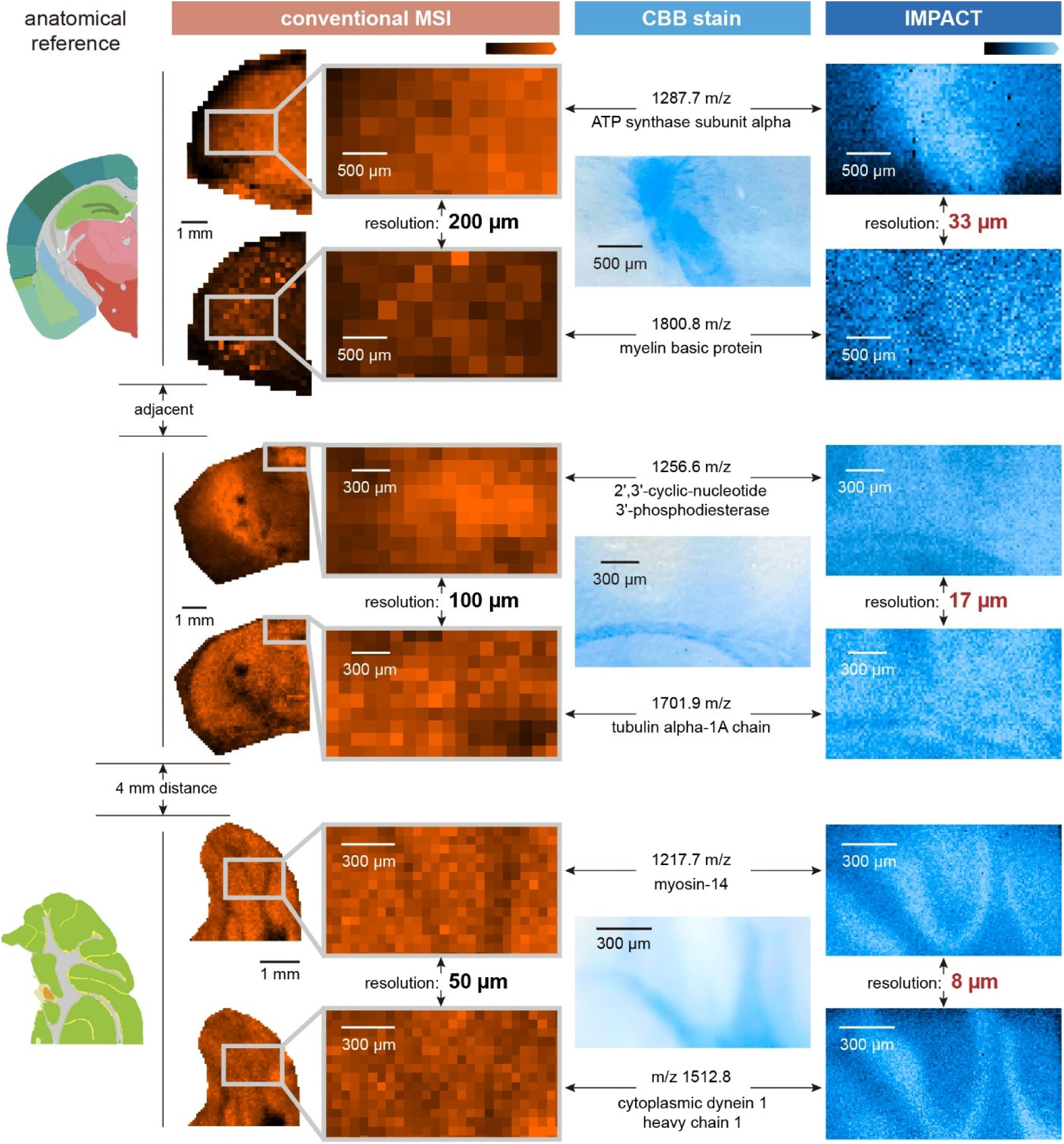
Images of representative peptides with protein identifications acquired with conventional MSI (10-µm-thick sections) and IMPACT (50-µm-thick sections) at varying spatial resolution settings, shown alongside corresponding anatomical reference (Allen Brain Atlas, left) and CBB stain (middle) of the same section used for IMPACT.

IMPACT-derived images demonstrated substantially improved clarity in protein distribution patterns compared to conventional MSI. The enhanced resolution not only provided superior image quality but also revealed fine-scale protein localization that closely matched anatomical references from both the mouse brain atlas and CBB staining of the same tissue section. This improvement enabled the visualization of protein distribution patterns associated with distinct cellular clusters and finely organized neuroanatomical regions. Notably, we observed distinct correlation patterns between protein distributions and CBB staining. Proteins such as ATP synthase subunit α exhibits a positive correlation that mirrored the CBB staining pattern. Proteins such as myelin basic protein exhibited inverse staining intensity relationships. Other proteins, such as myosin-14 and cytoplasmic dynein 1 heavy chain 1 showed localized enrichment along fiber tract boundaries as revealed by CBB stain. These differential correlation patterns underscore the critical need for proteome-wide, protein-specific spatial mapping within individual tissue sections. While conventional histological staining provides ensemble protein visualization, MSI-based spatial proteomics enables simultaneous resolution of multiple proteins’ distinct distribution patterns, offering unprecedented insight into their spatial organization and biological context.

### Enhanced spatial resolution facilitates integration with complementary spatial-omics approaches

The spatial distribution patterns of proteins identified through IMPACT can be validated using orthogonal protein imaging techniques such as immunofluorescence (IF). We acquired IF images of two representative proteins, i.e. the 26S proteasome non-ATPase regulatory subunit 13 (identified via peptide m/z 859.4), which plays a key role in ubiquitin-mediated protein degradation, and Thy-1 membrane glycoprotein (identified via peptide m/z 1162.6), a well-established marker for hematopoietic stem cells in nervous and immune systems. Notably, these proteins exist in low-abundance as revealed by LC-MS/MS-based proteomic analysis of an adjacent tissue section. IF patterns of these proteins in anatomically matched brain sections revealed consistent spatial patterns with IMPACT-MS results, showing higher abundance in fiber tract and brain stem regions compared to cerebrum (Figure 5). While IMPACT-MS still cannot match the resolution of optical microscopy, it provides the unique advantage of untargeted, multiplexed protein imaging without requiring specific probes or prior knowledge of proteins of interest.

**Figure 5.**
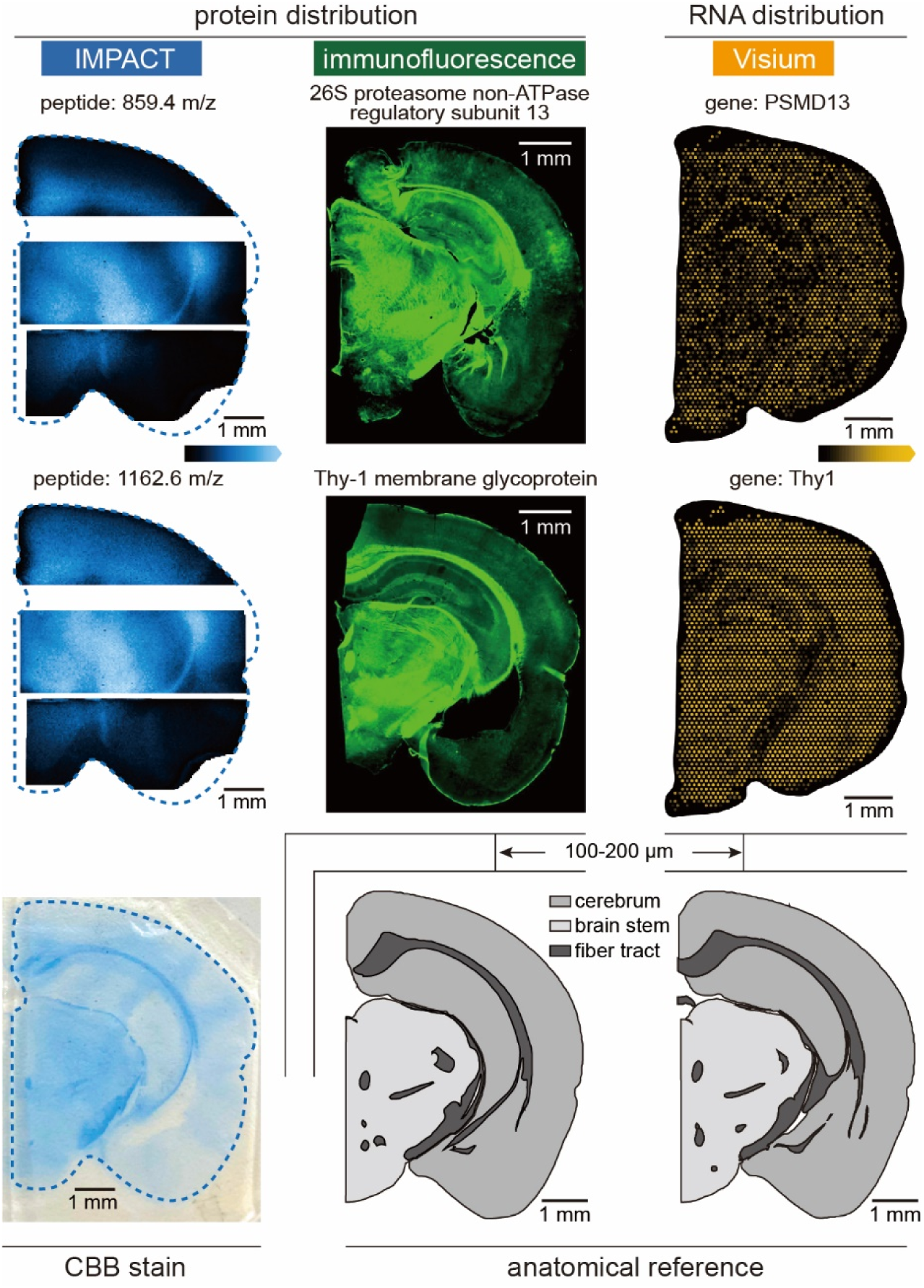
Comparative visualization of IMPACT-derived protein distributions (left), IF validation (middle), and Visium spatial transcriptomics (10x Genomics) from individual mouse brain sections. The bottom row shows CBB stain on the same IMPACT section and anatomical references from Allen Brain Atlas.

The relationship between protein localization and gene expression was further investigated by comparing IMPACT-MS results with spatial transcriptomics data acquired using Visium platform. For both PSMD13 (encoding 26S proteasome subunit 13) and Thy1 (encoding Thy-1 glycoprotein), we observed significant discordance between mRNA and protein distribution patterns (Figure 5). Compared with the protein images which exhibit more localized accumulation patterns, the corresponding transcripts showed relatively uniform expression across the tissue section with high abundance detected in cerebrum regions and lower abundance in fiber tracts. This observation was consistent across multiple gene-protein pairs, reinforcing that mRNA abundance alone cannot reliably predict protein localization. Such complementary nature of different spatial-omics approaches underscores the importance of direct protein spatial profiling, particularly for understanding post-transcriptional regulation and protein localization mechanisms in complex tissues.

## DISCUSSION

MS technique features high resolving power of signals and molecule identification based on unique masses. Accordingly, while MSI cannot match the sensitivity and spatial resolution of optical imaging techniques due to fundamental physical limitations, it provides unique advantages of accommodating higher imaging channels in a single scan and compatibility with untargeted discovery studies. Its value for spatial biology critically depends on two key parameters: (1) identification depth (number of detectable molecular species) which determines available imaging channels, and (2) spatial resolution which affects integration with complementary imaging modalities. These factors are particularly crucial for spatial proteomics, where the inability to amplify proteins (unlike nucleic acids) and the lack of signal amplification methods (without targeted probes) make efficient analyte utilization paramount.

While gel-assisted tissue expansion has proven successful in optical microscopy, its direct application to MSI presents unique challenges. The inherently lower sensitivity of mass spectrometry means that analyte dilution during 3D expansion, which is inconsequential for fluorescence microscopy, severely compromises MS signal intensity within each sampling pixel. This challenge is exacerbated for proteins, which have lower copy numbers and poorer ionization efficiency compared to small molecules. The IMPACT workflow addresses these limitations through vertical compression following expansion, which concentrates analytes vertically to increase surface density and reduces thickness-dependent electrical resistance to improve ionization efficiency, compensating for expansion-induced dilution while enhancing overall signal response. This approach simultaneously improves both spatial resolution and detection sensitivity, significantly increasing protein utilization rates and benefiting identification depth.

The IMPACT’s ability of analyzing thick tissue sections (>50 µm) allows expanding the range of viable specimens for biological and clinical research beyond conventional limits. First, it accommodates vibratome-prepared sections that better preserve native tissue architecture by eliminating freezing artifacts and avoiding embedding media that can interfere with MS signal, thereby enhancing data quality. Second, the technology facilitates analysis of challenging tissue morphologies that resist traditional sectioning approaches, including membrane-like structures (e.g., mesentery at mm scale, vascular walls at µm-mm scale, and retina at 100-300 µm thickness) and delicate structures such as cystic cavities. Third, IMPACT overcomes orientation-dependent limitations in pathological assessment, particularly for glandular tissues where abnormal cell progression may manifest along specific anatomical axes that would be obscured in unfavored sectioning planes. The method achieves these through thickness reduction that maintains spatial information and projecting three-dimensional spatial information into two-dimensional analyzable formats. Furthermore, IMPACT’s potential compatibility with Formalin-Fixed Paraffin-Embedded (FFPE) specimens promises its applications in retrospective clinical studies and archival tissue analysis.

The analytical value of mass spectrometry imaging depends on maintaining spatial fidelity throughout sample preparation. Artifactual diffusion of analytes or distortion of tissue during processing, even when accompanied by successful molecular detection, compromises the biological validity of resulting distribution patterns. Covalent protein anchoring within the gel matrix in IMPACT workflow ensures uniform tissue resizing while restricting molecular mobility to minimize diffusion. However, the Lys-gel linkages interfere with tryptic digestion by either limiting protease accessibility or generating undetectable cleavage products. This limitation is mitigated through use of complementary proteases such as chymotrypsin and Glu-C, which produce peptides between anchored lysine residues through their alternative cleavage specificities.

The enhanced spatial resolution and expanded protein coverage achieved by IMPACT enabled the spatial distributions of a broader range of proteins to be validated with IF images and compared with spatial transcriptomic data. While spatial transcriptomics is powerful for mapping mRNA distributions and cellular activities, it exhibits reported limitations in predicting protein abundance and localization^40, 41^. Our comparative analyses revealed spatial discordances between protein and corresponding mRNA distributions and highlighted the importance of direct proteomic measurement. The combination of high spatial resolution and deep proteome coverage enabled identification of fine-scale cellular heterogeneity that would be obscured in lower-resolution or less comprehensive datasets. These results underscore the complementary value of spatial proteomics and transcriptomics acquired with sufficient resolution and molecular coverage.

Further refinement of the IMPACT workflow should focus on two aspects: optimization of spatial fidelity preservation through improved gel-based resizing, and advancement of protein identification accuracy at the MSI stage. For spatial fidelity, current limitations arise from necessary specimen division prior to imaging, as standard glass slides cannot accommodate fully expanded tissues. This not only requires additional image realignment process that complicates data analysis workflow, but may also result in edge artifacts at gel division boundaries, manifesting as artificial signal elevation illustrated in Figure 3a. While pre-embedding specimen division could theoretically mitigate these artifacts, such an approach risks inducing tissue distortion due to loss of gel matrix stabilization during sectioning. Potential solutions include development of specialized instrumentation featuring enlarged ionization chambers and compatible sample supports and advanced computational algorithms for artifact correction and seamless image reconstruction.

The second major challenge lies in ensuring robust protein identification. Current MSI workflows predominantly rely on intact mass measurements, which become increasingly prone to false assignments as system complexity escalates, a particular concern in proteomic analyses where multiple peptides may share similar masses. While our implementation of adjacent-section LC-MS/MS validation provides fragmentation data for reliable identification for the peptide detected in the same data set, this approach cannot fully address the inherent challenges posed by natural peptide mass redundancies across the proteome and spatial heterogeneity between tissue sections. Direct integration of on-tissue tandem MS capabilities would represent the most rigorous solution, though substantial technical barriers remain regarding acquisition speed, fragmentation efficiency, and ionization sensitivity at high spatial resolution. Overcoming these limitations will require coordinated advances in both instrumentation and data acquisition methodologies to enable comprehensive, confident molecular identification while preserving the throughput necessary for practical imaging applications.

## CONCLUSION

We developed IMPACT, an MSI workflow that transforms spatial proteomics through integration of gel-based 3D expansion followed by vertical compression, in-situ in-gel proteolysis, MALDI-based high-resolution peptide imaging and LC-MS/MS-assisted protein identification. The horizontal expansion enhances spatial resolution to <8 μm, while the 100-fold thickness reduction of tissue section boosts sensitivity and enables analysis of thick (>50 μm) specimens. By increasing surface analyte density and implementing a hybrid-enzyme digestion strategy, IMPACT achieves a 3-fold improvement in protein identifications compared to conventional approaches. These advances collectively enable precise mapping of protein distributions to fine histological features and robust integration with orthogonal spatial-omics approaches. IMPACT thus provides a powerful platform for investigating spatial proteome organization in complex biological systems

## ACKNOWLEDGEMENTS

This work was supported by grants from National Health Commission of the People’s Republic of China (2023ZD0519900, 2023ZD0520300), National Natural Science Foundation of China (NSFC 22474005), Ministry of Science and Technology of the People’s Republic of China (2024YFA1306200) and Guangdong Basic and Applied Basic Research Foundation (2024A1515011065). The authors are grateful to Qingrong Chen, Dr. Lingxiao Chaihu and Dr. Mo Hu (Mass Spectrometry Core, Changping Laboratory) for help with MS operation, to Zhongqin Zhu (Shenzhen Bay Laboratory) for help with instrumentation, to Haotian Wang (Peking University) for helpful discussion, to National Center for Protein Sciences at Peking University for access to the digital slide scanner, and to Analytical Instrumentation Center, Peking University for access to the stylus profilometer.

